# Larger legume plants host a greater diversity of symbiotic nitrogen-fixing bacteria

**DOI:** 10.1101/246611

**Authors:** Russell Dinnage, Anna K. Simonsen, Marcel Cardillo, Peter H. Thrall, Luke G. Barrett, Suzanne M. Prober

## Abstract

A major goal in microbial ecology is to understand the factors that structure bacterial communities across space and time. For microbes that have symbiotic relationships with plants, an important factor that may influence their communities is host size or age, yet this has received little attention.

Using tree diameter size as a proxy for age, we quantified the diversity of rhizobia that associate with an endemic legume, *Acacia acuminata*, of variable size across a climate gradient in southwest Australia. We examined the 16S rRNA diversity (V1-V3 hypervariable region) of rhizobia at the taxonomic level and at higher sequence level diversity within taxonomic groups.

We identified 3 major taxonomic clades that associated with *Acacia acuminata:* Bradyrhizobiaceae, Rhizobiaceae, and Burkholderiaceae. Within these groups, we found extensive genetic variability, especially within Bradyrhizobiaceae. Using binomial multivariate statistical models that controlled for other factors that affect plant size and rhizobia community structure (climate and local soil characteristics), we determined that soil sampled at the base of larger *Acacia* trees was much more likely to contain a greater number of taxonomic clades and cryptic genetic variants within the Rhizobiaceae clade.

Despite strong influences of climate and highly heterogeneous soil conditions on rhizobial diversity, our results show that host tree size is a prominent factor in structuring nitrogen-fixing symbionts diversity across a large landscape. The identification of a positive relationship between plant host size and microbial diversity raise interesting questions about the role of plant host size in driving ecological processes that govern microbial community assembly. Specifically, our results suggest that hosts may modify the habitat of their surrounding soil to enhance growth (niche construction hypothesis) or that symbiotic microbes have large differences in dispersal capability. Our results also suggest that host plants may be analogous to ‘islands’, where larger legume hosts may accumulate diversity over time, through migration opportunities or in situ diversification. From a practical perspective, including plant size as an additional variable may assist sampling and analyses designs of future soil microbial studies.

## Introduction

Plants are highly dependent on mutualistic soil bacteria in the rhizosphere for growth and reproduction (Hayat et al. 2010; Berendsen, Pieterse, and Bakker 2012). Despite their functional importance, we are only beginning to understand the major ecological forces that structure mutualistic bacterial communities in the rhizosphere. Host plant size and age are ubiquitous, highly variable traits in most (if not all) natural plant communities (Hara 1988; Menges 2000). Through the simple act of growing, plants modify their surrounding environment, which in turn can have important top-down effects on microbial rhizosphere diversity (Philippot et al. 2013) by altering numerous ecological and evolutionary processes (i.e. selection, drift, diversification and dispersal) directly relevant to community assembly patterns (Vellend 2010; Nemergut et al. 2013). Among the key mutualistic microbes in the rhizosphere, symbiotic nitrogen fixing bacteria (i.e. rhizobia) are critical to ecosystem function due to their intimate associations with legumes and their ability to directly convert atmospheric nitrogen into a plant available nutrient (van der Heijden et al. 2006; Kahindi et al. 1997). Here, we evaluate the influence of variable plant size, within a keystone legume species, on the diversity and structure of symbiotic rhizobia communities.

Theoretically, host plant size and age have important impacts on soil microbe diversity. Through selection (i.e. niche-based processes), differences in diversity may be generated because hosts may actively or passively modify the rhizosphere as they grow and create more niches (i.e. “niche construction” (Odling-Smee et al. 2013)) (Meaden, Metcalf, and Koskella 2016; Marques et al. 2014; Micallef et al. 2009; Chaparro, Badri, and Vivanco 2014; Wagner et al. 2016). Alternatively, larger hosts may create more habitable space compared to smaller hosts, thus harbouring greater diversity associated with greater immigration or in situ diversification opportunities (i.e. bigger or smaller island in the context of Island Biogeography Theory or Neutral Theory) (Hubbell 2001; MacArthur and Wilson 2015). For example, larger hosts may harbour greater microbial diversity because their surrounding rhizosphere has higher absolute amounts of hospitable space for soil microbes. Host age, which is highly correlated with plant size, may critically affect rhizosphere diversity due to ecological drift (leading to loss of diversity) or accumulation of lineages with slower colonization rates, leading to an increase in diversity (Nemergut et al. 2013). For example, as legume hosts become older, they may accumulate larger rhizobia populations, making mutation and in situ diversification more likely.

The impact of plant size on microbial diversity through selection or dispersal can be either positive or negative, however diversification or ecological drift are generally predicted to show a positive relationship. Given that any of these community assembly processes, in isolation or in combination, may lead to either a positive or negative relationship between plant size and soil microbe diversity, the aim of this study is to evaluate the extent to which such relationships exist in natural mutualistic microbe populations. To gain further insight into the effects of plant size in the face of other important environmental drivers acting at either highly localized (i.e. edaphic conditions) or regional scales (i.e. climate), we sampled communities over a complex heterogeneous landscape. The interaction between legumes and rhizobia provide an ideal biological system to evaluate relationships between plant size and mutualistic microbial diversity because symbiotic rhizobia also exist as free-living bacteria in the soil and are not entirely dependent on their plant host to reproduce or disperse to other habitats. This potentially allows more dynamic community assembly processes as plants increase in age or size. Furthermore, the functional significance of rhizobia towards legume growth and reproduction is firmly established.

In this study, we examined the effect of host plant size on the diversity of rhizobia that originate from the surrounding soil of natural *Acacia acuminata* populations, a foundational species of the nationally endangered York gum (*Eucalyptus loxophleba*) ‐jam (*Acacia acuminata*) woodlands of the Western Australian wheatbelt (Prober, Standish, and Wiehl 2011). Using metagenomic 16S sequence data of symbiotic rhizobia isolated from bulk soils in vicinity of *A. acuminata* trees, we investigated the role of host size in influencing two levels of symbiotic rhizobia diversity: at large clade levels indicative of taxonomic levels, and at more cryptic genetic diversity levels within clade groups. We sampled rhizobia from the surrounding soil of *A. acuminata* host plants across the maximum climate gradient of the species range, capturing rhizobia diversity across a broad range of environments. Our analyses show that larger hosts are more likely to have a greater diversity of rhizobial taxa and a greater number of cryptic within-taxa genetic diversity, thus indicating the critical role that legume host size plays in shaping symbiotic rhizobia soil communities. We found host size is important for rhizobia diversity even after accounting for climate and soil related factors likely to be important for rhizobia diversity, at both local within-site, and regional between-site levels.

## Materials & Methods

### Site selection and soil sampling

*Acacia acuminata* is an endemic tall legume shrub or small tree restricted to south-western Australia. We sampled rhizobia from 24 sites where *A. acuminata* occurred. To select sample sites, we obtained *A. acuminata* occurrence records from the Atlas of Living Australia (ALA, www.ala.org.au) and associated temperature (mean annual temperature) and rainfall (mean annual rainfall) values. We chose sites to optimize three criteria: 1) maximize temperature and rainfall gradient; 2) minimize spatial autocorrelation in rainfall and temperature; and 3) no correlation between temperature and rainfall values. Based on the available occurrence records in ALA, our final combination of sites ranged in mean annual temperature from 15.4C to 20.6C, and mean annual rainfall from 287mm to 612mm. Temperature and rainfall were not correlated in our final site list (r=-0.1560238, p=0.47).

We sampled soil during the winter season (July) when the soil likely contains higher numbers of viable rhizobia due to higher seasonal rainfall in the region and higher levels of nitrogen-fixation between the host and its rhizobial partner (Monk, Pate, and Loneragan 1981). At each site, we selected three *A. acuminata* trees at least 50 meters from the road and at least 50 meters apart (where possible). We sampled three (5 cm diameter × 10 cm depth) soil cores within 50 cm of the base of each tree to obtain one pooled soil mixture for every sampled tree. We measured the circumference of each sampled tree and converted the measurement to diameter, as a proxy for plant age. We measured circumference at the base of each tree to accommodate comparison among different forms of *A. acuminata*, since this species can also occur in more shrub like forms. While some sampling was from these more shrub like forms, we refer to each sampled plant as a ‘tree’ throughout this paper. Soil was stored at 4C until further processing in the lab.

### Rhizobia isolation and DNA extraction

We isolated symbiotic rhizobia from field soil by harvesting legume nodules of *Acacia acuminata* grown in controlled growth chamber conditions inoculated with each field soil sample, where a field soil sample comprised a mixture of three soil cores collected from around the base of one *Acacia* tree. Wild collected *Acacia acuminata* seeds (‘typical form’, sourced from native seed collectors Nindethana Seed; <u>www.nindethana.net.au/</u>) were surface sterilized and pre-germinated on 1% agar plates for one week at room temperature (25C) in the dark. Six randomly selected *Acacia* seedling replicates were grown in each field soil sample, in a randomized blocked design in a growth chamber. For each pot, the field soil was sandwiched between two layers of autoclaved vermiculite and planted with *A. acuminata* seeds. Plants were watered weekly with 10 ml of ¼ strength McKnight solution and autoclaved water. Plants were grown for 4 weeks, allowing ~ 2 weeks for nodule formation. We sampled 5 nodules from each *Acacia* seedling, and restricted nodule sampling to below the first layer of autoclaved vermiculite to avoid any potential contaminants. Control pots containing only sterile vermiculite were randomized throughout the growth chamber (n=10), which had few or negligible nodules at harvest. Surface sterilized nodules (using commercial bleach) were crushed with sterile forceps and tissue lysate was cultured on Yeast-Mannitol agar. Rhizobia cultures were grown for 7-15 days at 30C in the dark. Isolates were replated twice on YMA agar media to obtain a single rhizobia isolate. Genomic DNA was extracted using MoBio kits (UltraClean Microbial Isolation kit), following the standard kit protocol.

In total, our sampled rhizobia collection consisted of ~1900 isolates, where ~30 rhizobia isolates were sampled from each *Acacia* tree (i.s. field soil sample), and ~90 rhizobia isolates were sampled from each site, representing an approximately equal sampling effort of rhizobia isolates for every *Acacia* tree and site.

### Sequencing

Our unit of metagenomic sequencing in this study was at the tree level (n=72 trees, 3 trees/site, 24 sites in total). We prepared pooled rhizobia DNA samples at the tree level by suspending freshly grown cells using a 1ul inoculation loop into 500 ul of sterile autoclaved distilled water, pooling ~25-30 isolates per sequencing sample. Pooled bacteria isolates were sent to AGRF for DNA extraction and 16S amplicon metagenomic sequencing, sequencing the v1-3 hypervariable region following standard AGRF amplification and library preparation protocols, producing 300 base-pair paired end reads from a single Miseq run.

### Raw sequence data processing

For each metagenomic sample (i.e. *Acacia* tree) paired reads were merged (using Flash), creating ~ 450bp long reads after Illumina adaptor trimming. In each sample, after discarding chimeras using Decipher and all singletons, a preliminary BLAST showed that the top hit for each read expectedly returned rhizobia taxa (Bradyrhizobiaceae, Rhizobiaceae, and Burkholderiaceae). To reduce the occurrence of sequence error in our final sequence dataset, we removed any reads that made up less than 0.5% of a sample. Assuming that most culture plates represented a single colony, the theoretical maximum of approximately 30 rhizobia isolates was expected per sample. After we applied our sequence filtering steps, the number of unique reads present in each sample was below 30 (between 2 and 18 reads). Out of the total cultured isolates examined in this study (approximately 1900), we obtained 95 unique 16S unique reads (after filter steps were applied) classified as rhizobia across all samples.

### Environmental variables

To examine relationships with our community data, we focused on three major classes of ecological and environmental factors: 1) climate; 2) physical and chemical soil characteristics; and 3) host size and host density. Mean annual precipitation and temperature were obtained from the Atlas of Living Australia (ALA: <u>www.ala.org.au</u>; also used for the preliminary site selection –see above), and are based on Bioclim (Fick and Hijmans 2017). For each soil sample, we measured 22 soil chemistry factors [see Table S1; analyses were carried out by CSBP (www.csbp-fertilisers.com.au)]. We measured the trunk diameter at the base of each tree that was sampled as a proxy of host size. We obtained estimates of site level host density using observation records available from ALA. We calculated site-level *Acacia acuminata* density using all occurrence record data for *Acacia acuminata* in South Western Australia found in the Atlas of Living Australia. We estimated *Acacia* density from point occurrence records using two-dimensional gaussian kernel estimation, implemented in the kde2d function of the MASS package in R. Density was estimated within a grid with a rough resolution of 0.37 degrees latitude-longitude within each cell, which is capable of capturing among site variation. Values for each site were then assigned as the density of the cell in which they fell. All factors, with the exception of climate data, were measured at the soil sample level (i.e. tree level).

### Data Analyses

The goal of our analyses was two-fold: firstly, to determine what environmental factors are associated with fine-scale rhizobia genetic diversity, as measured by the number of unique 16S reads and secondly, to determine whether richness responses to environmental factors depend on the major clade associated with a major taxonomic group. We first identified the major clades in our rhizobia sequence by inspecting the phylogeny. We used MUSCLE (Edgar 2004) to align all unique reads retained in our dataset and constructed a phylogeny in BEAST (Bouckaert et al. 2014). We identified 3 major taxonomic groups in our sample using BLAST (Madden 2013) (one with reads blasting to the Bradyrhizobiaceae, one with reads blasting to the Rhizobiaceae, and one blasting to the family Burkholderiaceae). Therefore, in total we delineated 3 major clades (Figure 1).

**Figure 1.**
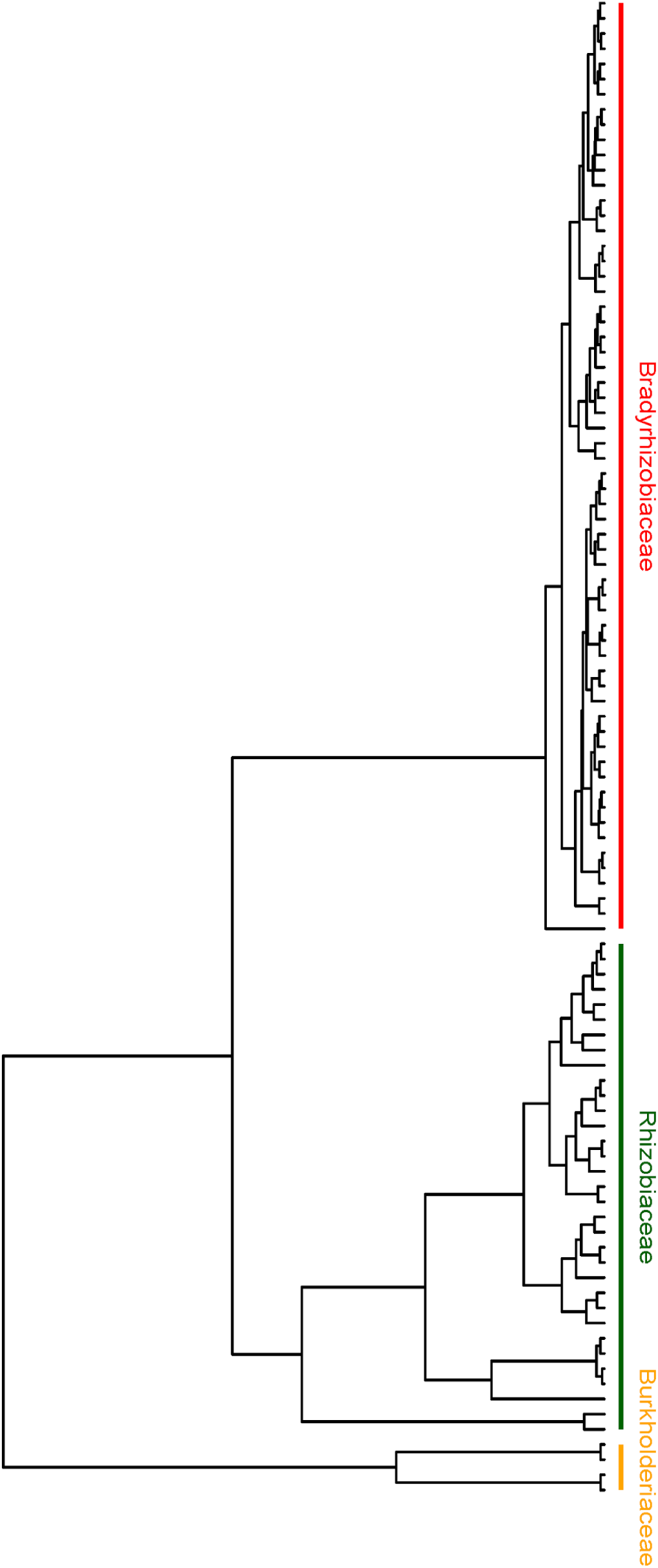
Bayesian phylogeny of all unique 16S reads detected in this study.

For soil (chemical and physical) characteristics, we collapsed the initial set of 22 variables into a smaller number using principal coordinates analyses. For our downstream analyses, we used the first four PC axes, which cumulatively explained ~70% of variation. When we inspected soil PC 1-4, we found no particular subset of soil factors that had particularly high loadings (Table S1, Figure S3).

In total, we analyzed 9 predictors in all subsequent statistical models (Mean Annual Temperature, Mean Annual Rainfall, Soil PC 1-4, Tree diameter, Latitude, *Acacia* Density). Given the large number of explanatory variables (relative to the available degrees of freedom), we used a model selection approach to determine which explained the most variation in our sequence diversity data. We determined that convergence and sample size was limited when we incorporated more than 4 variables, and so restricted model selection between a set of models which had no more than 6 environmental variables. A maximum of 4 environmental factors in each model gave us 10 degrees of freedom (excluding random effects; 4 * 2 clades interactions + 2 clade specific intercept terms). With 72 sample locations in total, this gives approximately 7 samples per degree of freedom, providing a reasonable power in our models, while avoiding overfitting and still providing the ability to statistically infer the main effects of the most influential factors. The form of the models is described below.

### Statistical Model

To model the 16S read data we used a multivariate model-based approach. The object of the analysis was to model the distribution of unique 16S reads with respect to a set of (at most) 6 environmental factors and the major clade they belonged to. We used a generalized linear mixed model (GLMM) with binomial errors to model the presence or absence of unique 16S reads in individual tree samples. Though our read data also included a measure of the abundance of each read in each sample, we did not model this data because it is not comparable between samples due to uncontrolled sources of variation in the total number of reads measured per identified read group. Specifically, preliminary DNA extraction trials on a subset of individual isolates showed large differences in total DNA yield, despite approximately equal cell inputs in the extraction protocol. Therefore we considered only the presence or absence of a read in a sample to be reasonably reliable data. Given this, our model estimates the effect of environment and clade on the probability of a read in an *Acacia* tree-level sample. An increased probability of reads in a tree-level sample is indicative of higher read diversity, and so this is a model of unique read diversity within the surrounding soil of an *Acacia* tree.

We included random effects for the tree, the site, and the unique read in all of our models to fully account for any non-independence due to these hierarchical factors. Fixed effects include the effect of the clade to which a read belonged, and the clade by environment interactions, encoding the effects of each environmental factor for each clade independently. The equation describing our full model structure is:

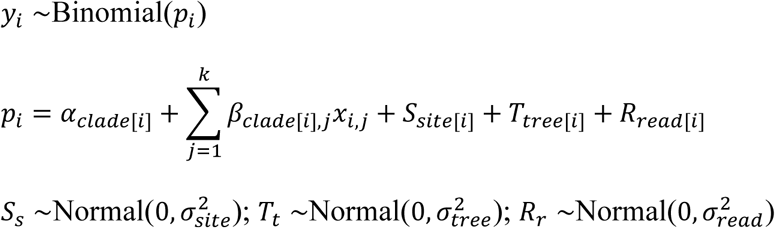

where *y_i_*is a response vector of ones and zeroes describing whether a particular read was found in a particular (tree-level) sample. *clade*[*i*], *site*[*i*], *tree*[*i*], and *read*[*i*]refer to the clade, site, tree, and read for record *i*. *α_c_*;is a fixed intercept for clade *c*, *β_c,j_*and is a fixed coefficient describing the effect of environmental variable *j*on reads reads that belong to clade *c*. *k*is the total number of environmental variables in the model, and ranges from 1 to 4, depending on the model. *S_s_*, *T_t_*, and *R_r_*refer to the random effects for site *s*, tree *t*, and read *r*, respectively. 
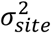
, 
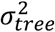
, and 
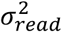
are the variance terms for the random effects of site, tree and read, respectively.

Models were fit using the R package `lme4` (Bates et al. 2015). As an example, the command to run one of the above models with 4 environmental variables was as follows:

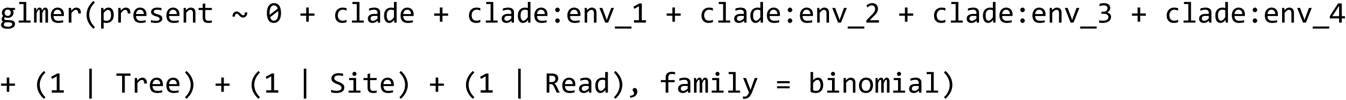

where `present` was a factor variable denoting presence or absence, `clade` is a factor variable denoting the clade, `env_i` is environmental variable `i, Tree` is a factor variable denoting the tree, `Site` a factor denoting the site, and `Read` a factor denoting the read.

We ran models for all possible combination of 4 environmental variables (for a total of 210 models). The model with the lowest AICc (Akaike Information Criterion with sample size correction) was retained as the top explanatory model and most subsequent analyses were based on it. AIC values were transformed to Akaike weights, providing conditional probabilities for each model (Burnham and Anderson 2003), which were used to estimate the relative importance of each environmental factor by summing the Akaike weights across all models containing each variable. Once a best model was chosen, we tested how well it conformed to model assumptions by examining its Dunn-Smyth residuals, which are designed to deal with non-gaussian integer response data (Dunn and Smyth 1996; Warton et al. 2017). We also tested its goodness-of-fit, by comparing the predicted read counts at individual Acacia tree samples with the observed counts. Predicted counts were calculated by summing the predicted probabilities of individual reads within each tree sample.

Subsequent models (see “Separating Within and Between Site Effects of Environment”), were based on the best model according to AICc.

### Separating Within and Between Site Effects of Environment

Our analysis was explicitly hierarchical, in that the data was measured within individual trees, which were found at different sites. Though we used hierarchical random effects in a mixed model to account for the non-independence induced by this sampling strategy, it is still interesting to ask how much of the effects we see can be attributed to within site differences amongst trees, versus between site differences. This information is lost in a simple linear mixed model, as it can only test for an overall effect (which therefore blends any within-site effect with any between-site effect). To examine within-site and between-site effects of the environment, we used a method known as contextual analysis (Snijders and Bosker 2011). In contextual analysis, the effects of a variable on the response is modelled as a regression of both the individual measured variable, and the mean of the variable for the group that the individual datapoint belongs to. Here, we modelled the presence or absence of a clade in a tree sample within a site as a function of the environment measured at the individual tree, as well as the mean environment for the site. With the exception of climate variables, all of our environmental variables were measured at the tree-level, and so we were able to decompose them into a site-level mean, and a tree-level deviation from the site mean. Mathematically, we created two new variables 
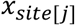
, and *Δx_j_* from each tree-level variable *x_i_* retained in the best model, such that:

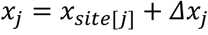

We included both the site-level mean 
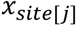
and the tree-level deviation from the site-level mean *Δx_j_* in a new model (Table 3). The goal of this model was to see whether any of the variables measured at the tree-level were actually driving compositional changes in rhizobia mainly at the site-level, in which case it would be more difficult to rule out the effect is not a result of local processes driven by the host tree, but possibly by a confounding site-level process.

**Table 1.** Importance of different factors in determining rhizobia community clade structure. Importance is calculated as the sum of Akaike weights for models containing the term. Excluded are any factors contained in all models (e.g. Clade, Intercept).

| Factor | Importance |
| --- | --- |
| Soil PC 1 | 0.97 |
| Annual Mean Rainfall | 0.97 |
| Host Tree Diameter | 0.80 |
| Soil PC 4 | 0.36 |
| Soil PC 2 | 0.34 |
| Latitude | 0.19 |
| Soil PC 5 | 0.14 |
| Annual Mean Temperature | 0.11 |
| Soil PC 3 | 0.07 |
| Acacia Density | 0.04 |

**Table 2.** Final ‘best’ generalized linear model (according to AICc), modelling the presence or absence of all unique 16S reads occurring in the metagenomic sample using clade specific diversity responses to environmental factors as predictors.

| Fixed Effects | Estimate | z value | P(>\|z\|) |
| --- | --- | --- | --- |
| Bradyrhizobiaceae (Intercept) | -2.868 | - | - |
| Rhizobiaceae (Intercept) | -4.543 | - | - |
| Bradyrhizobiaceae:Tree_diam | -0.0073 | -0.099 | 0.921 |
| Rhizobiaceae:Tree_diam | **0.514\*\*\*** | 4.066 | <0.0001 |
| Bradyrhizobiaceae:Env_ann_rain | **0.284\*\*\*** | 2.598 | 0.0094 |
| Rhizobiaceae:Env_ann_rain | **-0.536\*\*\*** | -2.642 | 0.0082 |
| Bradyrhizobiaceae:Soil_1 | -0.140 | -1.389 | 0.164 |
| Rhizobiaceae:Soil_1 | **0.622\*\*\*** | 3.860 | 0.00011 |
| Bradyrhizobiaceae:Soil_2 | 0.0144 | 0.149 | 0.881 |
| Rhizobiaceae:Soil_2 | **-0.356164\*\*** | -2.426 | 0.015 |
| Random Effects | Number of Levels | Estimated Variance | Standard Deviation |
| Site | 24 | 0.147 | 0.383 |
| Tree | 72 | 0.030 | 0.173 |
| Unique Read | 95 | 1.856 | 1.362 |
| *Note:* *p<0.1; **p<0.05; ***p<0.01 |  |  |  |

**Table 3.** Final ‘best’ generalized linear model, modelling the presence or absence of all unique 16S reads occurring in the metagenomic sample using clade specific diversity responses to environmental factors as predictors. This model simultaneously accounts for variation within sites and between site effects by incorporating predictor values representing site-level means (“_mean”), as well as individual Acacia tree-level deviations from the site-level means (i.e. within-site effects; “_dev”) for each factor (except rainfall, which as a climate factor, did not differ between trees within a site).

|  | Estimate | z value | P(>\|z\|) |
| --- | --- | --- | --- |
| Bradyrhizobiaceae (Intercept) | -2.90 | - | - |
| Rhizobiaceae (Intercept) | -3.28 | - | - |
| Bradyrhizobiaceae:Tree_diam_mean | 0.01928 | 0.081 | 0.94 |
| Rhizobiaceae:Tree_diam_mean | **1.77\*\*\*** | 3.982 | < 0.0001 |
| Bradyrhizobiaceae A:Tree_diam_dev | -0.01 | -0.118 | 0.906 |
| Rhizobiaceae:Tree_diam_dev | **0.59649\*\*\*** | 3.041 | 0.002 |
| Bradyrhizobiaceae A:Env_ann_rain | **0.328\*\*\*** | 2.648 | 0.00809 |
| Rhizobiaceae:Env_ann_rain | **-0.52012\*\*** | -2.166 | 0.030 |
| Bradyrhizobiaceae A:Soil_1_mean | -0.012 | -0.170 | 0.866 |
| Rhizobiaceae:Soil_1_mean | **0.492\*\*** | 2.082 | 0.037 |
| Bradyrhizobiaceae A:Soil_1_dev | 0.0135 | 0.082 | 0.935 |
| Rhizobiaceae:Soil_1_dev | **1.072\*\*\*** | 2.845 | 0.0044 |
| Bradyrhizobiaceae A:Soil_2_mean | -0.075 | -0.604 | 0.546 |
| Rhizobiaceae:Soil_2_mean | -0.33007 | -1.612 | 0.107 |
| Bradyrhizobiaceae A:Soil_2_dev | 0.154 | 0.828 | 0.408 |
| Rhizobiaceae:Soil_2_dev | **-1.104\*\*** | -2.374 | 0.0176 |
| Random Effects | Number of Levels | Estimated Variance | Standard Deviation |
| Site | 24 | 0.14 | 0.37 |
| Tree | 72 | 0.024 | 0.16 |
| Unique Read | 95 | 1.86 | 1.36 |
| Note: | *p<0.1; **p<0.05; ***p<0.01 |  |  |

### Interpreting Model Results

We interpreted the results of the best model according to AICc by examining its parameter estimates and by generating useful combinations and summaries of parameters through parametric bootstrapping.

Our model allowed us to estimate and evaluate the effects of each environmental factor on diversity of reads within each of our clades, but we were also interested in the overall diversity of reads. We were able to evaluate the effects of each environmental variable on read diversity by predicting how the number of unique reads was expected to change across environmental factors within each clade, and in total, using parametric bootstrapping to evaluate confidence intervals. We predicted data from our model for each set of bootstrapped coefficients on evenly spaced values of each environmental factor, within its observed range in our data, and then summarized that data to the predicted mean and confidence interval of counts before plotting them. We calculated the predicted count of each clade in a tree with a particular environmental value by multiplying the predicted probability of a read in a tree by the total number of unique reads (thus giving us the expected number of reads for that clade). We also noted that the predicted changes in differ ent clades would lead to different patterns of clade-level diversity for different values of the environmental variables. To explore this further, we calculated the Gini-Simpson diversity index (Jost 2006), from the predicted counts and plotted the mean and confidence interval for this against each environmental variable. The Gini-Simpson diversity index measured the diversity of types (i.e. clades) as the probability that two randomly drawn individuals from the community are of different types, and is at a maximum when all types are equally likely to be found in a community. Results were examined by plotting the predictions and their confidence intervals (Figures 2 and 3).

**Figure 2.**
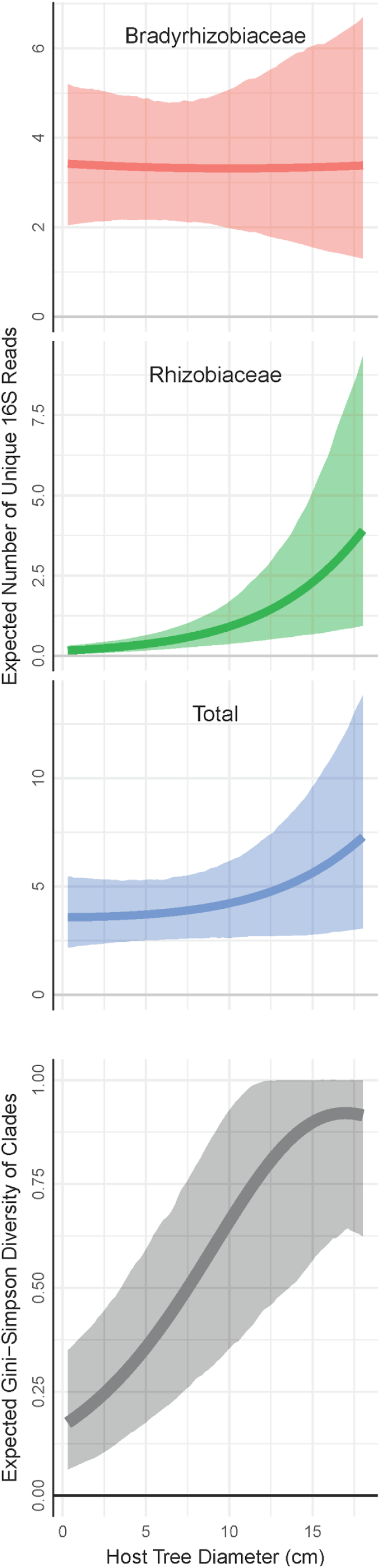
Total and clade level unique 16S read diversity responses to Acacia tree size Acacia tree diameter. Estimates are based on model predictions, controlling for other correlated effects (soil chemistry). 95% confidence bands are based on parametric boot bootstrapping. Bottom panel shows the predicted mean and 95% confidence interval for Gini-Simpson diversity of clades as a function of tree diameter. The Gini-Simpson metric measures the probability that two randomly drawn reads belong to different clades.

**Figure 3.**
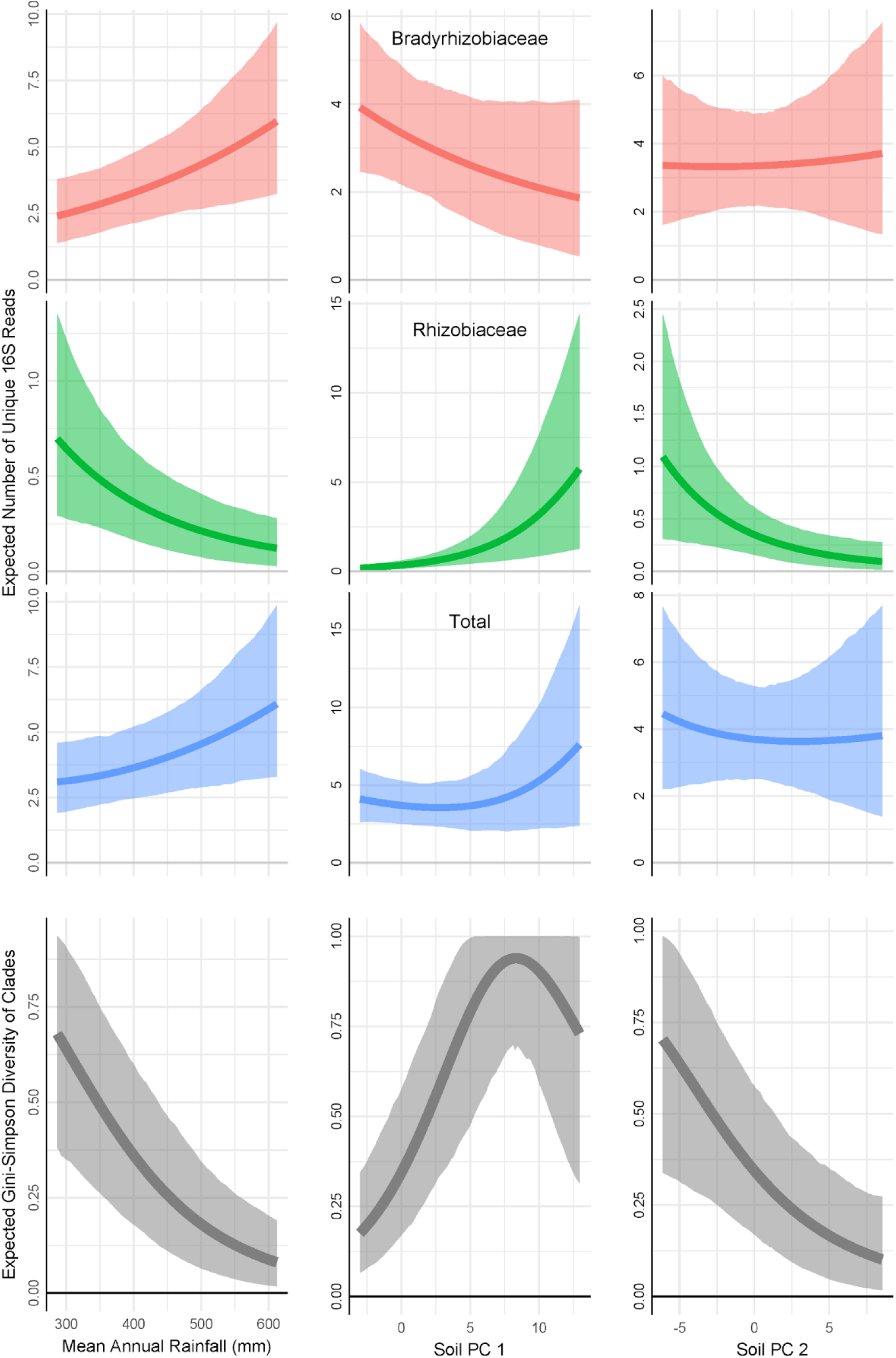
Total and clade level unique 16S read diversity responses to rainfall and soil chemistry. Estimates are based on model predictions, controlling for other effects. 95% confidence bands are based on parametric bootstrapping. Bottom panel shows the predicted mean and 95% confidence interval for Gini-Simpson diversity of clades as a function of the same factors. The Gini-Simpson metric measures the probability that two randomly drawn reads belong to different clades. Soil PC1 and Soil PC2 are the first two axes from a principal components analysis on 22 soil variables.

## Results

Out of the total input of ~1900 cultured isolates across all samples, we obtained 100 unique 16S reads classified as rhizobia after filtering sequence data which are commonly found in association with Australian acacias (Lafay and Burdon 1998; Hoque, Broadhurst, and Thrall 2011). All unique 16S read sequences could be classified into 3 distinct clades (Figure 1). Bradyrhizobiaceae clade had the most number of unique reads (62 unique reads), followed by Rhizobiaceae (33 unique reads), while *Burkholderia* was only represented by 7 unique reads. With the exception of Burkholderiaceae, at least one unique read from each major clade occurred in all soil samples (i.e. at every sampled acacia tree across all 24 sites), while *Burkholderia* was only detected in 2 soil samples across 2 sites. Due to the limited diversity and distribution of *Burkholderia* reads, we excluded this clade from our presented analyses, but inclusion or exclusion of this clade did not alter any inferences on Bradyrhizobiaceae and Rhizobiaceae.

The best model for predicting our sampled 16S rhizobia diversity, according to AIC, included the following environmental variables: Mean Annual Rainfall, Soil PC 1, Host Tree Diameter, and Soil PC 2.

Additional confidence on the explanatory power of retained environmental factors in the best model was also reflected in the weighted importance of each factor estimated across all evaluated models (although Soil PC 4 is slightly more important than Soil PC 2 across all models). More specifically, Mean Annual Rainfall, Soil PC 1 and Host tree size were weighted with high importance when summing across all models that contained them (0.80 or higher), and all other factors had considerably lower summed Akaike weights (0.36 or lower; Table 2). Abiotic factors, mean annual rainfall and soil PC 1 had the highest importance (Table 2, 0.97), *Acacia* tree diameter also had a high importance (0.80), indicating that plant size (in addition to abiotic factors) is a strong predictor of rhizobial diversity.

Though AICc can choose the best model out of the set of models tested, it cannot say how well that model fit the data in an absolute sense. We found that the best model fit the 16S read data fairly well when we examined its residuals and its predicted values. We used Dunn-Smyth residuals, which are designed to deal with non-gaussian integer responses (Dunn and Smyth 1996; Warton et al. 2017). There was no evidence of violation of the assumptions of the GLMM (Figure S1), with good a good fit to normality of errors, and no evidence of non-homogeneity of variance. Predicted read counts from the models, when summarised at individual tree samples, showed a strong relationship with the observed read counts (Pseudo-R^2^ = 0.85; figure S2).According to our best model, Annual Mean Rainfall, Soil PC 1, Host Tree Diameter, and Soil PC 2 predicted rhizobial diversity at two broad levels: 1) at the unique 16S read level, and 2) at the major clade level (i.e. taxonomic level). We describe these responses in greater detail below.

### Host tree diameter predicts rhizobia diversity within and among sites

We found a greater likelihood of observing more clades and unique 16S reads (Figure 2) from soil sampled at the base of larger *A. acuminata* trees. Furthermore, the likelihood of detecting a higher diversity of 16S read sequences within the Rhizobiaceae clade increased as tree size increased (Table 1; Figure 2). These analyses generally indicate that diversity among and within clades was higher in larger trees, where changes in diversity patterns were being driven entirely by the presence or absence of unique reads within Rhizobiaceae.

We further analyzed the possibility that Rhizobiaceae were more likely to occur at higher diversity in larger tree hosts within or among sites. In other words, was there evidence that higher diversity observed in larger trees was because some sites generally had larger *Acacia* trees, or was there evidence that even within sites of variable tree size, a higher diversity of rhizobia could still be observed at larger trees? As with the previous analyses, our results consistently show strong diversity responses in Rhizobiaceae, especially at the site level (Table 3). However, we also found a greater probability of observing higher Rhizobiaceae diversity (Table 3) for larger trees that occurred within a given site. In total, our analyses on the effects of tree size on rhizobia diversity show with confidence that Rhizobiaceae is much more likely to be present in larger trees, and this diversity sorting effect was even predictive for larger trees within sites when other highly localized factors (soil characters) are also included in the analyses.

### Rainfall predicts rhizobia clade diversity

We generally found strong patterns of rhizobia diversity associated with rainfall, with Bradyrhizobiaceae and Rhizobiaceae showing differential responses (Figure 2). We observed more clades at drier sites. When we examined the diversity patterns of unique reads within each clade, we found that Rhizobiaceae had a greater number of unique reads at sites with low rainfall. In contrast, our fitted model shows a greater number of unique Bradyrhizobiaceae reads at high rainfall. When examining the net effect of all unique reads, we found higher genetic diversity at wetter sites (Figure 2). In total, our rainfall results are similar to tree size in that Rhizobiaceae occurrences and diversity appear to be limited to more restricted conditions (i.e. drier sites), while Bradyrhizobiaceae occur along the entire rainfall gradient, and showing higher levels of diversity at wetter sites, the net effect translating into higher total unique read diversity at wetter sites.

### Soil characteristics predict rhizobia diversity within and among sites

We found strong patterns of diversity associated with soil characteristics. Similar to rainfall and host tree diameter responses, we found clade level diversity responses. However in contrast to rainfall and host tree diameter responses, we did not find any strong total read responses associated with soil PC1 and PC2. These results generally indicate a compositional shift in clades, but no general pattern of increase or decrease in diversity in response to soil characters. Generally, soil PC1 was quite weighted evenly by most chemical related factors variables, while PC2 was more uniquely weighted by physical factors compared to PC1 (soil texture and particle size; Figure S3). Further inspection of soil PC1 and PC2 indicate that PC1 is more strongly correlated with mean annual rainfall, while soil PC2 is more strongly associated with tree diameter (Table S3).

## Discussion

In this study, we evaluated the effect of plant size on the diversity of rhizobial symbiont communities across variable abiotic climate and soil conditions within a single dominant legume species. In summary, we found that soil at the base of larger *Acacia* trees hosted a higher diversity of rhizobia. Specifically, we found that larger trees had a higher Simpson diversity index at the taxonomic clade level as well as at the unique 16S read level. Out of the 3 major taxonomic clades we identified in this study (Fig 1) Bradyrhizobiaceae was equally likely to be present across all host plant sizes. Therefore the increase in the diversity was primarily driven by the presence or absence of Rhizobiaceae, which were much more likely to occur at the base of larger *Acacia* trees. Furthermore, when we examined the relationship between tree size and rhizobial diversity within each clade, we found a higher diversity of unique reads within Rhizobiaceae from soil found at larger trees, indicating that tree size also affected cryptic genetic variation. While accounting for other abiotic factors that also potentially covaried with host plant size, we found that climate (mean annual rainfall) and edaphic conditions were strongly associated with compositional shifts and that Rhizobiaceae in particular was highly responsive to changes in environmental conditions. This suggests that Rhizobiaceae generally has a much narrower environmental range compared to Bradyrhizobiaceae, but its response to tree size was distinct to other environmental factors.

Larger *Acacia* trees (on average) tend to be found at wet sites (Table S2), but even after incorporating rainfall into our analyses, tree size was still a strong determinant of rhizobia communities. In fact, our analyses showed that rainfall was negatively associated with Rhizobiacea diversity, such that the direct effect of rainfall and its indirect effect through its positive association with host size seem to be driving rhizobia diversity in opposing directions. We also showed that the tree size affect was as strong at the local within-site level as it was at the regional between-site level using a statistical method known as contextual analysis (Table 3), making it less likely that the effect is driven by a large-scale unmeasured confounding factor, and consistent with a mechanism driven by local-scale processes. We are not aware of many previous uses of this statistical method in ecology (but see Bradford et al. 2017), but a related method has been used in evolutionary biology – contextual selection analysis – designed to distinguish group selection from individual-level selection coefficients (Goodnight, Schwartz, and Stevens 1992). However, the method is well established in the social sciences (e.g. Davis, Spaeth, and Huson 1961; Enders and Tofighi 2007; Gelman et al. 2008; Snijders and Bosker 2011; Bell, Jones, and Fairbrother 2017), and would appear to have great potential in ecology, given that it allows modelling and testing of different effects of predictors at different scales. Rhizobial diversity patterns also responded strongly to local soil chemistry, in addition to larger scale factors acting at the regional site level (i.e. mean annual rainfall) (Table 3).

The finding that larger legumes have higher symbiotic microbe diversity in the surrounding soil has several implications for the community assembly processes of mutualistic microbes, which we discuss below.

Our results are consistent with the hypothesis that as trees grow in size or age, they modify the habitat of their surrounding soil, generating niches that can support higher rhizobial diversity. This may therefore constitute an empirical example of niche construction, where an organism modifies its existing habitat and subsequently imposes selective forces on other interacting organisms (e.g. bacteria in the rhizosphere) at the community level (Odling-Smee et al. 2013). Modification of a plant’s own rhizosphere is by definition a highly local process and previous studies have shown that rhizobial population dynamics are also highly localized in that rhizobial population density is much higher in the immediate vicinity of legume roots, while soil meters away contains significantly fewer rhizobia (Parker, Malek, and Parker 2006). Consistently, we find that the influence of tree size on rhizobial community assembly is also highly localized, since our results indicate that rhizobia (and Rhizobiaceae in particular), are strongly responsive to within-site variation in tree size.

How *Acacia acuminata* or other plants actively or passively modify their rhizosphere requires further investigation. However, our data indicates a shift in soil chemical composition as a potential modification pathway, since we found that soil PC2 strongly predicts *Acacia* tree size (Table S2), and that one of the unique characteristics of PC2 is a much higher loading of ammonia (NH3) and iron (Figure S3). Prober *et al*. (2011) also found high levels of ammonia and iron at the base of *Acacia acuminata* sampled from soil in the same woodland habitats (i.e.York-gum jam woodlands). Given that iron has been identified as a critical and limiting element for nitrogen-fixation (Brear, Day, and Smith 2013), and that ammonia is a known product of biological nitrogen fixation reaction that can be utilized by legumes, our results raise the possibility that legumes are modifying chemical aspects of their rhizosphere to enhance nitrogen-fixation function and increase recruitment success of nearby legume seedlings. However, our study also demonstrates that other unmeasured changes associated with increasing tree size are likely to be taking place, given that tree size is still strongly explanatory even after controlling for soil conditions in our analyses.

Apart from niche-driven differences, our results could also be explained by differences in dispersal capacity or competitive ability among rhizobial clades or taxa (Bissett et al. 2010). Given the wide distribution of Bradyrhizobiaceae, it is possible that Bradyrhizobiaceae has a much higher dispersal, establishment or competition capability and thus may be more important for new *Acacia* seedling recruitment, which is also consistent with the observation that *Bradyrhizobium* is more frequently found across Australia compared to *Rhizobium* (Lafay and Burdon 1998). If Rhizobiaceae has a slower dispersal or recruitment rate, it may only begin to become common in older plants, which have had more time to accumulate rhizobia lineages.

The explanations above assert that rhizobial taxa exhibit different characteristics or particular habitat preferences. In contrast, species equivalency is one of the key assumptions of metacommunity dynamics in the context of neutral theory (Hubbell 2001), and is of increasing interest in microbial ecology (Dumbrell et al. 2010). Generally, we find the assumption of equivalency among our identified rhizobial clades unlikely. More specifically, under the assumption of neutrality, and its prediction of high diversity with larger area, we would expect a relationship with total unique reads and host size, but no relationship between clade diversity and host size, since differential clade level responses imply non-neutral, higher level phylogenetic organization driving community assembly patterns. However, we found both unique read and clade level diversity associated with host size and strong clade specific responses to different abiotic conditions (climate and soil characteristics), suggesting that clades are not equivalent (although equivalency may hold within clades). More specifically, Rhizobiaceae appears to occur over a much narrower range of edaphic and climate conditions. Previous studies also find that differences in abiotic growth conditions is variable among rhizobial taxa and strains (Li et al. 2011; Han et al. 2009; Thrall, Bever, and Slattery 2008; Vuong, Thrall, and Barrett 2017), which together imply that species equivalency is unlikely in natural rhizobia populations. Together, these results build a stronger case for niche driven differences that will benefit from further experimental work.

Few studies thus far have sought to explicitly examine the impact of plant size on microbial diversity among natural natural plant populations (though see Meaden, Metcalf, and Koskella 2016), despite the recognition that variation in plant size (reflected by either the productivity or age of the habitat) is a prominent and highly dynamic feature of plant communities (i.e. natural forest stands are always a mixture of different age stages). Previous manipulative transplant work has shown that soil microbe communities change during the course of plant development (Wagner et al. 2016; Philippot et al. 2013), implying that plant age is likely to be generally important for microbes. At the same time, other manipulative studies have shown that inoculating legumes with a higher diversity of rhizobial strains leads to decreased host plant performance (Simonsen, Chow, and Stinchcombe 2014; Barrett et al. 2015). If diversity of rhizobia does reduce plant growth (perhaps due to interference effects) this opens the interesting possibility that the increase in diversity with larger plants could lead to a negative feedback effect that will eventually limit the ability of an *Acacia* tree to grow larger.

To our knowledge this is the first ecological study to demonstrate a positive relationship between legume plant size and naturally occurring symbiotic rhizobial diversity. In addition to demonstrating a positive relationship between plant size and soil symbiont diversity, our study also demonstrates that the effect of tree size on microbial diversity is sufficiently strong that it can even be detected across a large complex landscape of high climate and soil variability. Together, these studies contribute to a growing body of evidence that plant size is important in structuring the diversity of soil microbe communities, and highlight the potentially important role of host size and age in modifying ecological processes in the rhizosphere. Finally, these studies provide additional considerations that can be informative for future sampling design wishing to measure patterns of microbial diversity in natural populations.

## Acknowledgments

We would like to thank Cathryn O’Sullivan and Alan Richardson for helpful comments on the manuscript. This research was funded in part by an Ignition Grant from the Centre for Biodiversity Analysis, Canberra, Australia.

## Author’s Contributions

AS created the sampling design and experimental idea, in consultation with SP. RD and AS collected the data, analysed the data, and led the writing of the manuscript. RD generated the figures. All authors contributed critically to the drafts and gave final approval for publication.

## Supplementary Information

### Supplementary Figures

**Figure S1.**
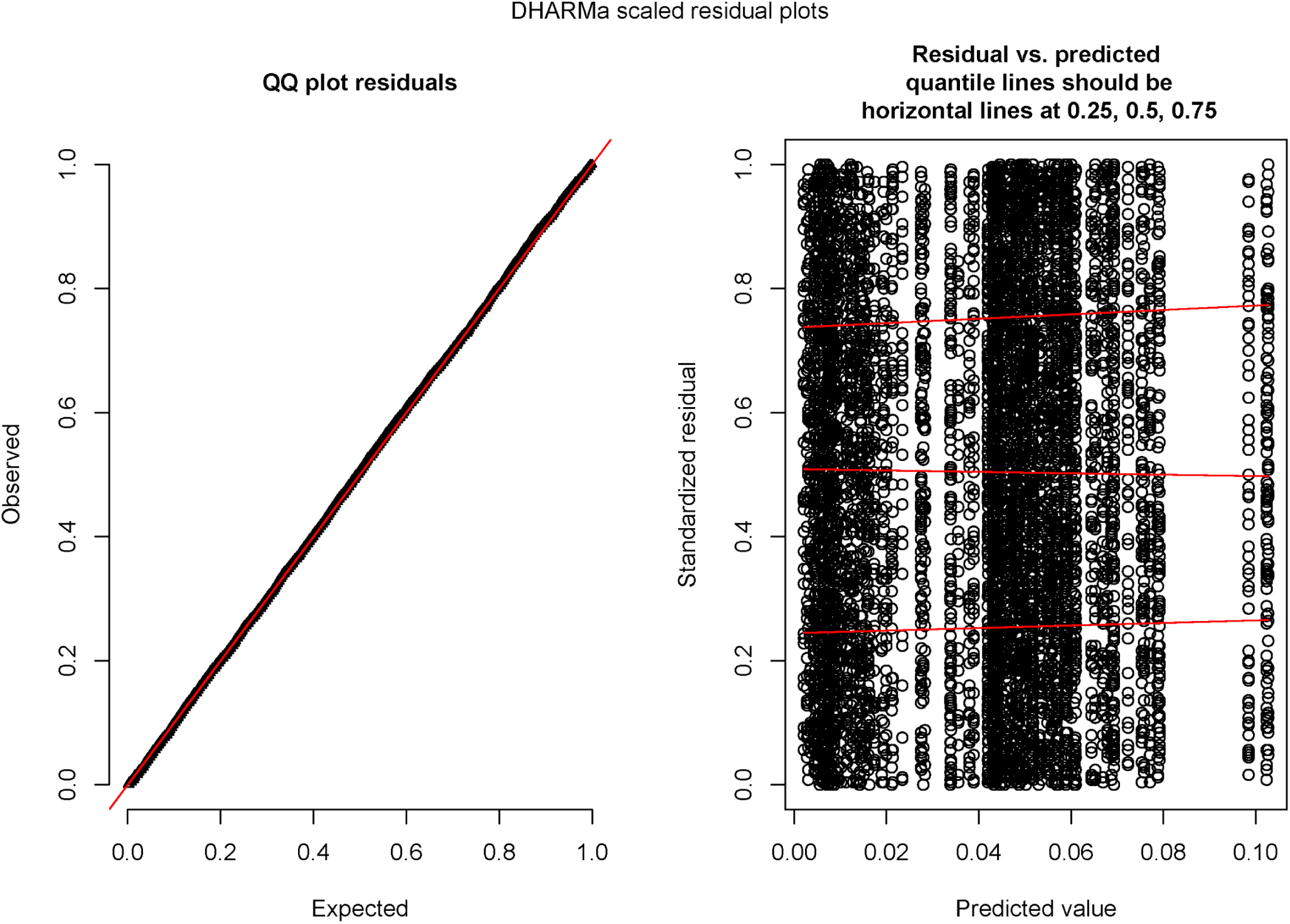
Dunn-Smyth Residuals from best model. Left panel shows quantile-quantile plot, showing model residuals match assumptions of normality nearly perfectly. Right plot shows that there is no evidence of non-homogeneity of variance in the residuals with respect to the predicted values.

**Figure S2.**
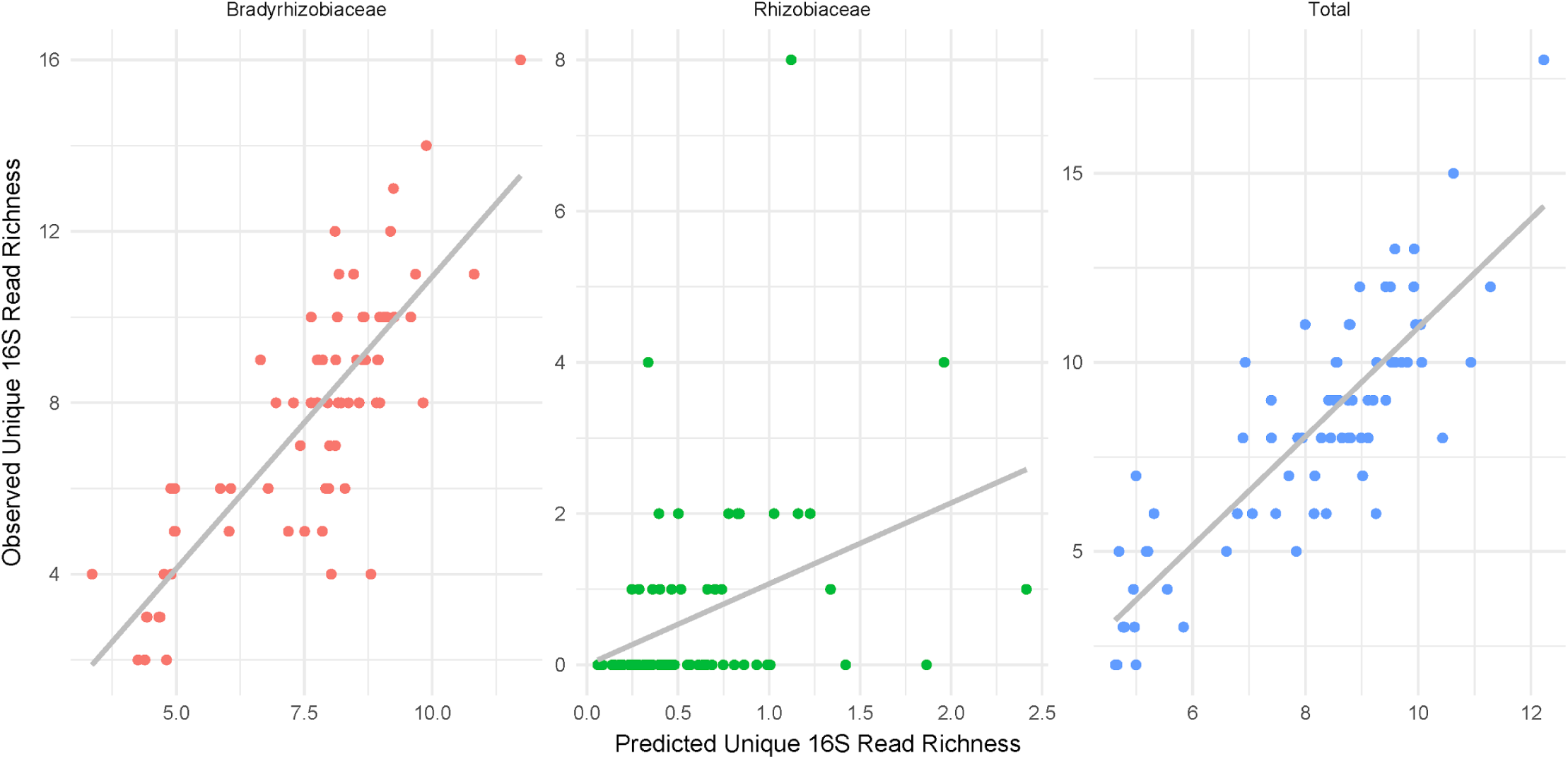
Goodness-of-Fit for best model. Total Pseudo-R^2^ = 0.85. Panels show predicted vs. observed read counts within samples from individual Acacia trees for each of the two clades and the total.

**Figure S3.**
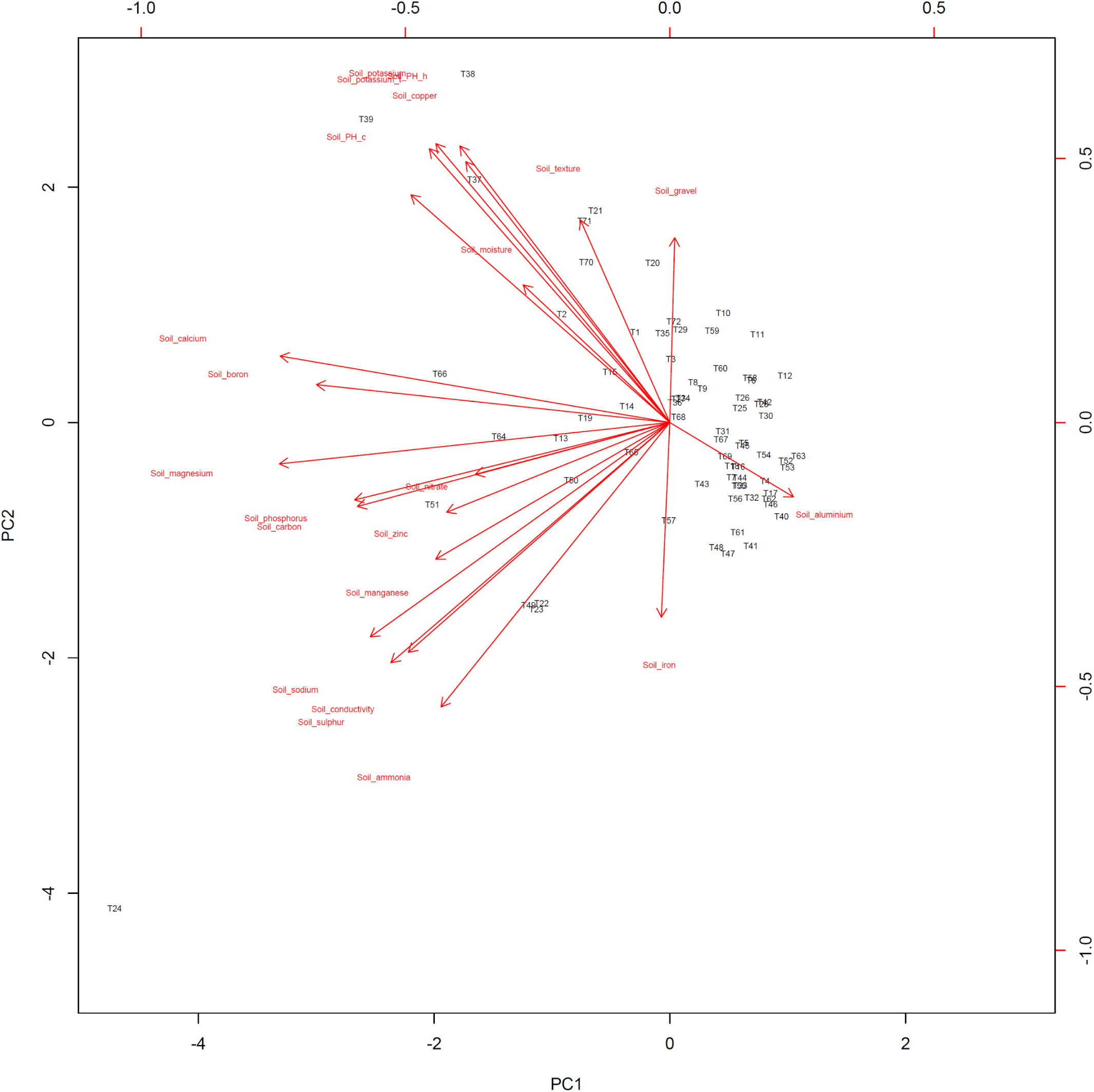
Soil PC 1 and Soil PC2 biplot.

### Supplementary Tables

**Table S1.** Soil PC loadings.

|  | PC1 | PC2 | PC3 | PC4 | PC5 | PC6 | PC7 | PC8 | PC9 | PC10 | PC11 |
| --- | --- | --- | --- | --- | --- | --- | --- | --- | --- | --- | --- |
| Soil_moisture | 0.125 | -0.156 | -0.176 | 0.297 | -0.390 | 0.151 | 0.148 | -0.562 | 0.290 | -0.307 | 0.057 |
| Soil_gravel | -0.004 | -0.210 | 0.212 | 0.430 | 0.077 | 0.033 | -0.567 | 0.089 | -0.191 | -0.340 | 0.167 |
| Soil_texture | 0.076 | -0.230 | 0.189 | 0.444 | -0.034 | 0.300 | -0.130 | -0.148 | -0.178 | 0.524 | -0.287 |
| Soil_ammonia | 0.196 | 0.323 | 0.094 | -0.176 | 0.003 | 0.099 | -0.227 | -0.229 | -0.042 | -0.135 | -0.145 |
| Soil_nitrate | 0.166 | 0.059 | 0.232 | 0.300 | 0.393 | -0.415 | 0.163 | -0.074 | -0.098 | -0.301 | -0.036 |
| Soil_phosphorus | 0.270 | 0.088 | 0.237 | -0.041 | 0.165 | 0.002 | -0.132 | 0.044 | 0.376 | 0.156 | -0.561 |
| Soil_potassium | 0.200 | -0.317 | 0.168 | -0.167 | -0.218 | -0.179 | 0.061 | 0.196 | -0.117 | -0.043 | -0.082 |
| Soil_sulphur | 0.239 | 0.273 | -0.199 | 0.176 | -0.131 | -0.076 | 0.013 | 0.216 | -0.127 | 0.113 | -0.017 |
| Soil_carbon | 0.267 | 0.095 | 0.298 | -0.073 | 0.141 | 0.051 | 0.137 | -0.094 | 0.353 | -0.030 | 0.143 |
| Soil_conductivity | 0.224 | 0.261 | -0.268 | 0.198 | -0.153 | -0.114 | -0.007 | 0.174 | -0.057 | 0.035 | -0.007 |
| Soil_PH_c | 0.221 | -0.259 | -0.252 | 0.032 | 0.258 | 0.081 | 0.038 | 0.029 | 0.034 | 0.172 | 0.309 |
| Soil_PH_h | 0.180 | -0.314 | -0.217 | -0.128 | 0.263 | 0.170 | 0.046 | -0.071 | 0.064 | 0.233 | 0.123 |
| Soil_copper | 0.175 | -0.296 | 0.032 | -0.154 | -0.393 | -0.251 | 0.005 | -0.109 | -0.130 | -0.021 | -0.149 |
| Soil_iron | 0.007 | 0.221 | 0.355 | -0.071 | -0.047 | 0.332 | 0.405 | -0.220 | -0.518 | 0.009 | 0.062 |
| Soil_manganese | 0.200 | 0.155 | 0.059 | -0.165 | -0.046 | -0.415 | -0.333 | -0.427 | -0.194 | 0.382 | 0.313 |
| Soil_zinc | 0.191 | 0.102 | 0.203 | -0.247 | -0.215 | 0.421 | -0.361 | 0.207 | 0.127 | -0.112 | 0.293 |
| Soil_aluminium | -0.106 | 0.085 | 0.394 | 0.307 | -0.282 | -0.167 | 0.214 | 0.188 | 0.364 | 0.298 | 0.362 |
| Soil_calcium | 0.333 | -0.075 | 0.034 | -0.039 | 0.211 | -0.036 | 0.055 | -0.127 | 0.080 | -0.080 | 0.164 |
| Soil_magnesium | 0.334 | 0.047 | -0.112 | 0.099 | -0.087 | 0.125 | 0.090 | 0.008 | 0.002 | -0.138 | -0.136 |
| Soil_potassium_t | 0.206 | -0.311 | 0.169 | -0.177 | -0.198 | -0.139 | 0.085 | 0.212 | -0.097 | -0.069 | -0.065 |
| Soil_sodium | 0.256 | 0.244 | -0.243 | 0.164 | -0.157 | -0.008 | -0.015 | 0.147 | -0.050 | 0.048 | -0.002 |
| Soil_boron | 0.302 | -0.043 | 0.043 | 0.077 | 0.125 | 0.171 | 0.221 | 0.246 | -0.206 | -0.039 | 0.146 |
|  | PC12 | PC13 | PC14 | PC15 | PC16 | PC17 | PC18 | PC19 | PC20 | PC21 | PC22 |
| Soil_moisture | -0.015 | 0.188 | -0.043 | 0.205 | -0.108 | -0.173 | 0.013 | -0.155 | 0.019 | -0.019 | 0.011 |
| Soil_gravel | 0.182 | 0.274 | 0.152 | -0.207 | 0.179 | 0.025 | -0.048 | -0.021 | -0.037 | 0.000 | -0.001 |
| Soil_texture | -0.070 | -0.297 | -0.251 | -0.040 | -0.146 | 0.030 | 0.077 | -0.012 | 0.053 | -0.027 | -0.015 |
| Soil_ammonia | 0.142 | -0.189 | 0.403 | -0.288 | -0.561 | -0.107 | 0.182 | 0.017 | -0.015 | -0.018 | 0.018 |
| Soil_nitrate | -0.456 | -0.223 | 0.010 | 0.184 | -0.170 | 0.046 | -0.147 | -0.127 | -0.056 | 0.006 | -0.038 |
| Soil_phosphorus | -0.052 | 0.381 | 0.175 | 0.218 | 0.227 | -0.216 | -0.008 | 0.051 | -0.081 | 0.028 | 0.006 |
| Soil_potassium | -0.015 | 0.175 | -0.157 | -0.070 | -0.273 | -0.028 | -0.071 | -0.050 | 0.087 | 0.156 | 0.696 |
| Soil_sulphur | -0.052 | 0.118 | 0.174 | 0.037 | 0.102 | 0.153 | 0.220 | -0.486 | 0.554 | -0.158 | 0.019 |
| Soil_carbon | 0.226 | -0.146 | -0.317 | -0.479 | 0.184 | -0.048 | -0.303 | -0.234 | 0.189 | -0.083 | -0.037 |
| Soil_conductivity | -0.108 | 0.069 | -0.213 | -0.226 | 0.014 | -0.185 | -0.003 | 0.126 | -0.467 | -0.549 | 0.130 |
| Soil_PH_c | -0.293 | 0.057 | 0.226 | -0.169 | -0.038 | -0.448 | -0.060 | 0.356 | 0.343 | 0.008 | -0.015 |
| Soil_PH_h | -0.063 | 0.163 | 0.282 | -0.083 | -0.113 | 0.381 | -0.157 | -0.406 | -0.398 | -0.066 | 0.011 |
| Soil_copper | -0.094 | -0.421 | 0.361 | -0.157 | 0.468 | -0.089 | 0.003 | -0.056 | -0.114 | 0.001 | -0.068 |
| Soil_iron | -0.185 | 0.345 | 0.102 | -0.121 | 0.192 | -0.030 | -0.023 | 0.048 | -0.061 | 0.004 | 0.015 |
| Soil_manganese | 0.140 | 0.136 | -0.116 | 0.262 | 0.012 | 0.011 | -0.177 | 0.060 | 0.022 | -0.012 | -0.023 |
| Soil_zinc | -0.448 | -0.189 | -0.103 | 0.286 | 0.024 | 0.034 | -0.041 | -0.067 | -0.043 | -0.044 | 0.007 |
| Soil_aluminium | 0.024 | 0.022 | 0.349 | -0.020 | -0.131 | 0.109 | 0.087 | 0.126 | -0.119 | 0.027 | 0.019 |
| Soil_calcium | 0.030 | -0.014 | -0.189 | -0.034 | 0.190 | 0.235 | 0.764 | 0.183 | -0.097 | 0.108 | 0.089 |
| Soil_magnesium | 0.135 | -0.064 | 0.111 | 0.096 | 0.028 | 0.561 | -0.343 | 0.528 | 0.176 | -0.083 | 0.034 |
| Soil_potassium_t | 0.013 | 0.255 | -0.174 | -0.041 | -0.310 | 0.026 | 0.084 | 0.032 | 0.050 | -0.151 | -0.667 |
| Soil_sodium | -0.067 | 0.047 | -0.120 | -0.180 | -0.011 | -0.030 | -0.116 | -0.038 | -0.204 | 0.769 | -0.187 |
| Soil_boron | 0.537 | -0.218 | 0.092 | 0.438 | -0.003 | -0.330 | -0.025 | -0.104 | -0.152 | -0.001 | 0.002 |

**Table S2.** Tree diameter, as predicted by other important predictors identified during model selection analyses.

|  | Estimate | t value | P(>\|t\|) |
| --- | --- | --- | --- |
| Intercept | 0.47 | - | - |
| Mean Annual Rainfall | **0.010\*** | 2.786 | 0.011 |
| Soil_PC1 | -0.16 | -1.296 | 0.20 |
| Soil_PC2 | **0.49\*\*** | 3.138 | 0.0038 |
| Random Effects | Number of Levels | Estimated Variance | Standard Deviation |
| Site | 24 | 0.68 | 0.83 |

*Note:* \*p<0.1; \*\*p<0.05; \*\*\*p<0.01

**Table S3.**
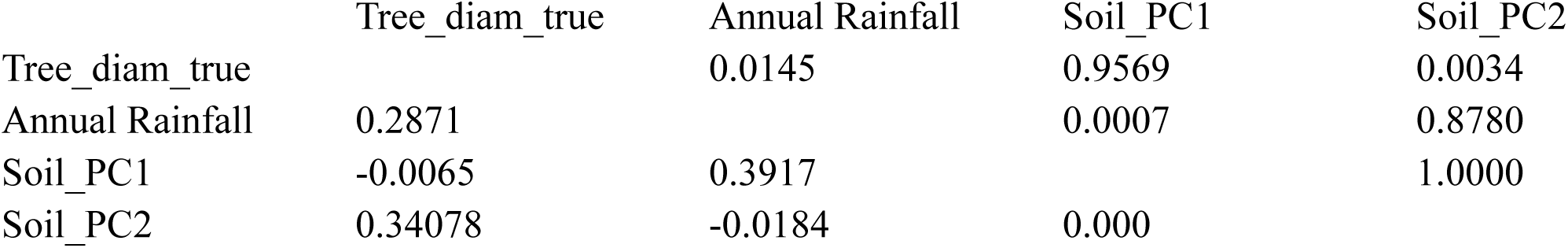
Correlation matrix among variables retained in best model (based on AIC). Bottom diagonal are spearman correlation coefficients and upper diagonals are p-values.

|  | Tree_diam_true | Annual Rainfall | Soil_PC1 | Soil_PC2 |
| --- | --- | --- | --- | --- |
| Tree_diam_true |  | 0.0145 | 0.9569 | 0.0034 |
| Annual Rainfall | 0.2871 |  | 0.0007 | 0.8780 |
| Soil_PC1 | -0.0065 | 0.3917 |  | 1.0000 |
| Soil_PC2 | 0.34078 | -0.0184 | 0.000 |  |

## References

Barrett, Luke G., James D. Bever, Andrew Bissett, and Peter H. Thrall. 2015. “Partner Diversity and Identity Impacts on Plant Productivity in Acacia–rhizobial Interactions.” The Journal of Ecology 103 (1):130–42.

Bates, Douglas, Martin Mächler, Ben Bolker, and Steve Walker. 2015. “Fitting Linear Mixed-Effects Models Using lme4.” Journal of Statistical Software, Articles 67 (1):1–48.

Bell, Andrew, Kelvyn Jones, and Malcolm Fairbrother. 2017. “Understanding and Misunderstanding Group Mean Centering: A Commentary on Kelley et Al.’s Dangerous Practice.” Quality & Quantity, November. Springer Netherlands, 1–6.

Berendsen, Roeland L., Corné M. J. Pieterse, and Peter A. H. M. Bakker. 2012. “The Rhizosphere Microbiome and Plant Health.” Trends in Plant Science 17 (8):478–86.

Bissett, A., A. E. Richardson, G. Baker, S. Wakelin, and P. H. Thrall. 2010. “Life History Determines Biogeographical Patterns of Soil Bacterial Communities over Multiple Spatial Scales.” Molecular Ecology 19 (19):4315–27.

Bouckaert, Remco, Joseph Heled, Denise Kühnert, Tim Vaughan, Chieh-Hsi Wu, Dong Xie, Marc A. Suchard, Andrew Rambaut, and Alexei J. Drummond. 2014. “BEAST 2: A Software Platform for Bayesian Evolutionary Analysis.” PLoS Computational Biology 10 (4):e1003537.

Bradford, Mark A., G. F. Ciska Veen, Anne Bonis, Ella M. Bradford, Aimee T. Classen, J. Hans C. Cornelissen, Thomas W. Crowther, et al. 2017. “A Test of the Hierarchical Model of Litter Decomposition.” Nature Ecology & Evolution 1 (12):1836–45.

Brear, Ella M., David A. Day, and Penelope M. C. Smith. 2013. “Iron: An Essential Micronutrient for the Legume-Rhizobium Symbiosis.” Frontiers in Plant Science 4 (September):359.

Burnham, Kenneth P., and David R. Anderson. 2003. Model Selection and Multimodel Inference: A Practical Information-Theoretic Approach. Springer Science & Business Media.

Chaparro, Jacqueline M., Dayakar V. Badri, and Jorge M. Vivanco. 2014. “Rhizosphere Microbiome Assemblage Is Affected by Plant Development.” The ISME Journal 8 (4):790–803.

Davis, James A., Joe L. Spaeth, and Carolyn Huson. 1961. “A Technique for Analyzing the Effects of Group Composition.” American Sociological Review 26 (2). [American Sociological Association, Sage Publications, Inc.]:215–25.

Dumbrell, Alex J., Michaela Nelson, Thorunn Helgason, Calvin Dytham, and Alastair H. Fitter. 2010. “Relative Roles of Niche and Neutral Processes in Structuring a Soil Microbial Community.” The ISME Journal 4 (3):337–45.

Dunn, Peter K., and Gordon K. Smyth. 1996. “Randomized Quantile Residuals.” Journal of Computational and Graphical Statistics: A Joint Publication of American Statistical Association, Institute of Mathematical Statistics, Interface Foundation of North America 5 (3). [American Statistical Association, Taylor & Francis, Ltd., Institute of Mathematical Statistics, Interface Foundation of America]:236–44.

Edgar, Robert C. 2004. “MUSCLE: Multiple Sequence Alignment with High Accuracy and High Throughput.” Nucleic Acids Research 32 (5):1792–97.

Enders, Craig K., and Davood Tofighi. 2007. “Centering Predictor Variables in Cross-Sectional Multilevel Models: A New Look at an Old Issue.” Psychological Methods 12 (2):121–38.

Fick, Stephen E., and Robert J. Hijmans. 2017. “WorldClim 2: New 1-Km Spatial Resolution Climate Surfaces for Global Land Areas.” International Journal of Climatology 37 (12). John Wiley & Sons, Ltd:4302–15.

Gelman, Andrew, Boris Shor, Joseph Bafumi, and David Park. 2008. “Rich State, Poor State, Red State, Blue State: What’s the Matter with Connecticut?” Quarterly Journal of Political Science 2 (4). Now Publishers:345–67.

Goodnight, Charles J., James M. Schwartz, and Lori Stevens. 1992. “Contextual Analysis of Models of Group Selection, Soft Selection, Hard Selection, and the Evolution of Altruism.” The American Naturalist 140 (5):743–61.

Han, Li Li, En Tao Wang, Tian Xu Han, Jie Liu, Xin Hua Sui, Wen Feng Chen, and Wen Xin Chen. 2009. “Unique Community Structure and Biogeography of Soybean Rhizobia in the Saline-Alkaline Soils of Xinjiang, China.” Plant and Soil 324 (1-2). Springer Netherlands:291–305.

Hara, T. 1988. “Dynamics of Size Structure in Plant Populations.” Trends in Ecology & Evolution 3 (6):129–33.

Hayat, Rifat, Safdar Ali, Ummay Amara, Rabia Khalid, and Iftikhar Ahmed. 2010. “Soil Beneficial Bacteria and Their Role in Plant Growth Promotion: A Review.” Annals of Microbiology 60 (4). Springer-Verlag:579–98.

Heijden, Marcel G. A. van der, Roy Bakker, Joost Verwaal, Tanja R. Scheublin, Matthy Rutten, Richard van Logtestijn, and Christian Staehelin. 2006. “Symbiotic Bacteria as a Determinant of Plant Community Structure and Plant Productivity in Dune Grassland.” FEMS Microbiology Ecology 56 (2):178–87.

Hoque, Mohammad S., Linda M. Broadhurst, and Peter H. Thrall. 2011. “Genetic Characterization of Root-Nodule Bacteria Associated with Acacia Salicina and A. Stenophylla (Mimosaceae) across South-Eastern Australia.” International Journal of Systematic and Evolutionary Microbiology 61(Pt 2):299–309.

Hubbell, Stephen P.. 2001. The Unified Neutral Theory of Biodiversity and Biogeography. Princeton University Press.

Jost, Lou. 2006. “Entropy and Diversity.” Oikos 113 (2). Blackwell Publishing Ltd:363–75.

Kahindi, J. H. P., P. Woomer, T. George, F. M. de Souza Moreira, N. K. Karanja, and K. E. Giller. 1997. “Agricultural Intensification, Soil Biodiversity and Ecosystem Function in the Tropics: The Role of Nitrogen-Fixing Bacteria.” Applied Soil Ecology: A Section of Agriculture, Ecosystems & Environment 6 (1):55–76.

Lafay, B., and J. J. Burdon. 1998. “Molecular Diversity of Rhizobia Occurring on Native Shrubby Legumes in Southeastern Australia.” Applied and Environmental Microbiology 64 (10):3989–97.

Li, Qin Qin, En Tao Wang, Yun Zeng Zhang, Yan Ming Zhang, Chang Fu Tian, Xin Hua Sui, Wen Feng Chen, and Wen Xin Chen. 2011. “Diversity and Biogeography of Rhizobia Isolated from Root Nodules of Glycine Max Grown in Hebei Province, China.” Microbial Ecology 61 (4):917–31.

MacArthur, Robert H., and Edward O. Wilson. 2015. Theory of Island Biogeography. Princeton University Press.

Madden, Thomas. 2013. The BLAST Sequence Analysis Tool. National Center for Biotechnology Information (US).

Marques, Joana M., Thais F. da Silva, Renata E. Vollu, Arie F. Blank, Guo-Chun Ding, Lucy Seldin, and Kornelia Smalla. 2014. “Plant Age and Genotype Affect the Bacterial Community Composition in the Tuber Rhizosphere of Field-Grown Sweet Potato Plants.” FEMS Microbiology Ecology 88 (2):424–35.

Meaden, S., C. J. E. Metcalf, and B. Koskella. 2016. “The Effects of Host Age and Spatial Location on Bacterial Community Composition in the English Oak Tree (Quercus Robur).” Environmental Microbiology Reports, April. https://doi.org/10.1111/1758-2229.12418.

Menges, E. S. 2000. “Population Viability Analyses in Plants: Challenges and Opportunities.” Trends in Ecology & Evolution 15 (2):51–56.

Micallef, Shirley A., Sheridon Channer, Michael P. Shiaris, and Adán Colón-Carmona. 2009. “Plant Age and Genotype Impact the Progression of Bacterial Community Succession in the Arabidopsis Rhizosphere.” Plant Signaling & Behavior 4 (8):777–80.

Monk, D., J. S. Pate, and W. A. Loneragan. 1981. “Biology of Acacia Pulchella R.br. With Special Reference to Symbiotic Nitrogen Fixation.” Australian Journal of Botany 29 (5). CSIRO PUBLISHING:579–92.

Nemergut, Diana R., Steven K. Schmidt, Tadashi Fukami, Sean P. O’Neill, Teresa M. Bilinski, Lee F. Stanish, Joseph E. Knelman, et al. 2013. “Patterns and Processes of Microbial Community Assembly.” Microbiology and Molecular Biology Reviews: MMBR 77 (3):342–56.

Odling-Smee, John, Douglas H. Erwin, Eric P. Palkovacs, Marcus W. Feldman, and Kevin N. Laland. 2013. “Niche Construction Theory: A Practical Guide for Ecologists.” The Quarterly Review of Biology 88 (1):4–28.

Parker, Matthew A., Wanda Malek, and Ingrid M. Parker. 2006. “Growth of an Invasive Legume Is Symbiont Limited in Newly Occupied Habitats.” Diversity and Distributions 12 (5). Blackwell Publishing Ltd:563–71.

Philippot, Laurent, Jos M. Raaijmakers, Philippe Lemanceau, and Wim H. van der Putten. 2013. “Going back to the Roots: The Microbial Ecology of the Rhizosphere.” Nature Reviews. Microbiology 11 (11):789–99.

Prober, Suzanne M., Rachel J. Standish, and Georg Wiehl. 2011. “After the Fence: Vegetation and Topsoil Condition in Grazed, Fenced and Benchmark Eucalypt Woodlands of Fragmented Agricultural Landscapes.” Australian Journal of Botany 59 (4). CSIRO PUBLISHING:369–81.

Simonsen, Anna K., Theresa Chow, and John R. Stinchcombe. 2014. “Reduced Plant Competition among Kin Can Be Explained by Jensen’s Inequality.” Ecology and Evolution 4 (23):4454–66.

Snijders, Tom A. B., and Roel J. Bosker. 2011. Multilevel Analysis: An Introduction to Basic and Advanced Multilevel Modeling. SAGE.

Thrall, Peter H., James D. Bever, and Jo F. Slattery. 2008. “Rhizobial Mediation of Acacia Adaptation to Soil Salinity: Evidence of Underlying Trade-Offs and Tests of Expected Patterns.” The Journal of Ecology 96 (4). Blackwell Publishing Ltd:746–55.

Vellend, Mark. 2010. “Conceptual Synthesis in Community Ecology.” The Quarterly Review of Biology 85 (2):183–206.

Vuong, Holly B., Peter H. Thrall, and Luke G. Barrett. 2017. “Host Species and Environmental Variation Can Influence Rhizobial Community Composition.” The Journal of Ecology 105 (2):540–48.

Wagner, Maggie R., Derek S. Lundberg, Tijana G. Del Rio, Susannah G. Tringe, Jeffery L. Dangl, and Thomas Mitchell-Olds. 2016. “Host Genotype and Age Shape the Leaf and Root Microbiomes of a Wild Perennial Plant.” Nature Communications 7 (July):12151.

Warton, David I., Jakub Stoklosa, Gurutzeta Guillera-Arroita, Darryl I. MacKenzie, and Alan H. Welsh. 2017. “Graphical Diagnostics for Occupancy Models with Imperfect Detection.” Methods in Ecology and Evolution / British Ecological Society 8 (4):408–19.

